# Genome-Wide Association Studies and Genomic Selection for Grain Protein Content Stability in a Nested Association Mapping Population of Spring Wheat

**DOI:** 10.1101/2021.04.15.440064

**Authors:** Karansher S. Sandhu, Paul D. Mihalyov, Megan J. Lewien, Michael O. Pumphrey, Arron H. Carter

## Abstract

Grain protein content (GPC) is controlled by complex genetic systems and their interactions, and is an important quality determinant for hard spring wheat as it has a positive effect on bread and pasta quality. GPC is variable among genotypes and strongly influenced by environment. Thus, understanding the genetic control of wheat GPC and identifying genotypes with improved stability is an important breeding goal. The objectives of this research were to identify genetic backgrounds with less variation for GPC across environments and identify quantitative trait loci (QTLs) controlling the stability of GPC. A spring wheat nested association mapping (NAM) population of 650 recombinant inbred lines (RIL) derived from 26 diverse founder parents crossed to one common parent, ‘Berkut’, was phenotyped over three years of field trials (2014-2016). Genomic selection models were developed and compared based on prediction of GPC and GPC stability. After observing variable genetic control of GPC within the NAM population, seven RIL families displaying reduced marker-by-environment interaction were selected based on a stability index derived from Finlay-Wilkinson regression. A genome-wide association study identified seven significant QTLs for GPC stability with a Bonferroni-adjusted *P* value <0.05. This study also demonstrated that genome-wide prediction of GPC with ridge regression best linear unbiased estimates reached up to *r* = 0.69. Genomic selection can be used to apply selection pressure for GPC and improve genetic gain for GPC.

## Introduction

Grain protein content (GPC) is a high-priority determinant of end-use quality for most cereals (Shewry and Halford 2002), including pasta (*Triticum turgidum* L.) and bread wheat (*Triticum aestivum* L.) where higher GPC is preferred. The large dependence on wheat, rice (*Oryza sativa* L.), and maize (*Zea mays* L.) as primary sources of carbohydrates results in their consumption meeting 80% of dietary protein requirements for humans (Shewry 2007). Compared to other agricultural commodities, cereal grains contain a relatively lower concentration of protein. In a screening of 12,600 lines from the USDA world wheat collection, GPC varied from 7% to 22%, with the genetic component accounting for only a third of the variation (Vogel et al. 1976). Breeding efforts to improve GPC have been difficult due to strong environmental influences and the high variability of GPC across years and locations (Löffler and Busch 1982), combined with variable economic value-based trade-offs between starch and protein yields in grains (DePauw et al. 2007).

Phenotypic plasticity refers to the flexibility for exhibiting changes to the different environments for a particular trait (Kroon et al. 2005). Improved understanding of genotype by environment interactions could provide new strategies for breeding crop varieties that remain stable for performance between changing environments (Kusmec et al. 2017). One of the key challenges to identifying genes controlling environmental stability lies in the quantification of stability as a trait (Kliebenstein et al. 2002). There are various approaches for measuring environmental stability including the use of variance components of individuals across environments (represented by coefficient of variation), the comparison of mean responses of genotypes to the overall mean of individuals in the trial (calculated as coefficient of regression on environment index) (Lin et al. 1986), and expected change in performance of genotype as function of environment effect (Finlay and Wilkinson 1963). Compared to unstructured genotype by environment interaction models, Finlay Wilkinson (FW) regression fits every level of genotype and environment and reveals genotype performance across environments (Lian 2016). GPC stability can potentially help select stable genotypes across increasingly unpredictable environments (Ordas et al. 2008). Various biparental and association mapping studies identified QTLs controlling GPC, but none of them focus on the stability of GPC (Blanco et al. 1996; Huang et al. 2010).

Nested association mapping (NAM) is a multi-parental design with higher allelic variation than bi-parental populations and stronger statistical power than association mapping populations, combining the advantages of each approach (Zhu et al. 2008). A universal parent is crossed to multiple genotypes, followed by inbreeding to make a combination of both full-sib and half-sib recombinant inbred lines (RILs) (Yu and Buckler 2006). NAM population have proven their success in mapping complex traits in barley (*Hordeum vulgare* L.) (Saade et al. 2016), maize (Gage et al. 2020), rice (Fragoso et al. 2017), and arabidopsis (*Arabidopsis thaliana*) (Li et al. 2011). The NAM population design mirrors the structure of several breeding programs where a few superior, diverse, or exotic lines are crossed with elite breeding lines during the pre-breeding pipelines. The cross of exotic germplasm with elite breeding lines for population development aids in normalizing genetic background and selecting segregating alleles in favor of the adapted parent alleles (Blanc et al. 2006). The resolution and power of NAM populations allow for the assessment of complex traits like GPC stability in structured germplasm via genome-wide association studies (GWAS).

Joint linkage association mapping is applied in multi-parental mapping populations where QTL terms are nested within families (Mcmullen et al. 2009) instead of testing marker effects across the families as in GWAS (Würschum et al. 2012). Larger allelic classes and more balanced allele frequencies increase power to detect QTLs with small effects. NAM combines better resolution by targeting historical recombination events in parents and also includes more causative events that are likely to segregate in biparental progenies (Tian et al. 2011). Thus, NAM is an effective tool for identifying major and minor QTLs associated with a particular trait (Korte & Ashley 2013).

In plant breeding, accumulation of minor QTLs is a major constraint given the dozens of loci and small population sizes. Hence, genomic selection (GS) is used to sum the effects of genome-wide markers to predict genomic estimated breeding values (GEBVs) (Meuwissen et al. 2001). In GS, a training population is genotyped and phenotyped for the traits of interest to estimate the genetic effect of each marker. The estimated genetic effect of each marker is then used to predict the GEBVs of a related population of breeding lines that is genotyped but has not been phenotyped. The selection of suitable genotypes among the test population is performed based on the predicted favorable allele GEBVs for each trait (Goddard et al. 2010). The success of GS depends upon the prediction accuracy of the GS model, which is measured as the correlation between estimated breeding values and the observed phenotypic values of the selected population (Crossa et al. 2014). GS can translate to higher genetic gains by reducing the number of progenies and cycles needed and also by improving selection intensity and reducing cycle time (Larkin et al. 2019; Robertsen et al. 2019).

Several studies have reported the application of GS for wheat yield and disease resistance (Sun et al. 2017; Sandhu et al. 2021a), but not thoroughly for quality traits, especially GPC and GPC stability. GS has been tested in soft winter wheat for end-use quality, where Heffner et al. (2011) found that end-use quality and processing traits are more predictive than grain yield. The prediction accuracy for flour yield and flour protein was 0.56 and 0.39, respectively, using a ridge regression model. GS accuracy for different agronomic traits and their stability was predicted in a winter wheat population of 273 lines and it was observed that GS accuracy varied from 0.33 to 0.67 for yield with yield stability having higher accuracy (Huang et al. 2016). Although several studies have focused on genomic selection for yield stability, none of them have investigated GPC stability. The main objectives of this study were to (1) detect marker-traits associations for GPC stability and GPC; and (2) identify ability of GS models for predicting GPC and GPC stability under cross and independent validations.

## Materials and Methods

### Plant material and trait measurement

Thirty-two spring wheat accessions from the USDA-ARS National Small Grains Collection were chosen as parental lines for the creation of the NAM population. These parents were crossed with the common cultivar ‘Berkut’ (‘Irena’/’Babax’//’Pastor’; released in 2002) to create 32 half-sib families (Blake et al. 2019). Berkut was included in this study because it is a broadly adapted photoperiod insensitive and semi-dwarf cultivar developed by the International Maize and Wheat Improvement Center (CIMMYT), Mexico.

Twenty-six families with genotype data provided by Kansas State University (Jordan et al. 2018) were then selected, and 25 random RILs from each family (650 total RIL; named NAM_650_) were planted between 2014 and 2016 at Spillman Agronomy Farm in Pullman, WA, under rainfed conditions (Sandhu et al. 2021b). A modified augmented design was used in each trial with three check cultivars (Berkut, ‘Thatcher’, and ‘McNeal’ (Lanning et al. 1994)). Planting was completed on 5^th^ May 2014, 8^th^ April 2015, and 10^th^ April 2016 based on field conditions. The agronomic practices, including nitrogen fertilizer goals, were uniform in each environment. Percentage of protein content in the grain was measured using a Perten DA 7000 NIR analyzer (Perkin Elmer, Sweden).

### Statistical analysis

Best linear unbiased estimates (BLUEs) were calculated with the augmented complete block design (ACBD) in the R program for individual environments (R Development Core Team. 2013; Rodríguezet al. 2018) using the model

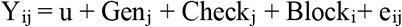

where Y_ij_ is the grain protein content of an individual line; µ is the mean effect; Block_i_ is the fixed effect of the i^th^ block, Gen_j_ represents the fixed effect of unreplicated genotypes; Check_j_ is the effect of the replicated checks within each block; and e_ij_ is the standard normal error. All the effects were considered fixed in BLUE calculations.

The significance of differences in GPC were analyzed across years and between families using analysis of variance (ANOVA). Standard deviation and coefficient of variation were calculated within the different families to identify those with lower variation for GPC across years. The GPC variation within families in each environment and across years was used to select families that had less variation for GPC. Broad sense heritability for GPC was obtained as

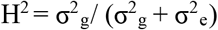

where σ^2^_g_ and σ^2^_e_ are the genotypic and error variance components, respectively.

### Stability analysis

The stability of each RIL was assessed using a Finlay Wilkinson (FW) program implemented in the FW package in R (Lian et al. 2016). The FW package jointly estimates the parameters of the FWR equation

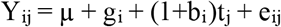

where Y_ij_ is the GPC of the i^th^ RIL from the j^th^ environment, µ is the mean effect, t_j_ is the main effect of the j^th^ environment, g_i_ is the main effect of the i^th^ RIL, (1+b_i_) is the change in expected performance of the i^th^ RIL per unit change in the environmental effect (Stability index), and e_ij_ is an error term assumed to be independent and identical distributed with mean zero and variance σ_e_^2^. All parameters are treated as random effects with distributions: **g** ∼ N(0, **A**σ^2^_g_), **b** ∼ N(0, **A**σ^2^_b_), and **t** ∼ N(0, **T**σ^2^_t_), where **A** is an N x N kinship matrix for all the RILs calculated from the complete set of genetic markers, and **T** is an M X M Pearson variance-covariance matrix, describing the relationship of phenotypic values among M environments.

The stability of each RIL was calculated using GPC values for a selected 175 RILs (NAM_175_) in each environment as dependent variables. The FW package returned the environmental stable genotypic effect (g_i_), environmental effect (t_j_), and the stability index of each RIL (b_i_). The stability index for each RIL provided an idea about plasticity across the environments (Kusmec et al. 2017). A stability index of 1 and −1 means that genotype is highly plastic and responds according to environmental changes whereas a stability index of 0 suggests the genotype performs stable under different environments. Loci associated with GPC stability were identified by GWAS of the stability index absolute values. The parents for the selected 175 RILs were ‘Dharwar Dry’, ‘PI210945’, ‘CItr15144’, ‘PI92569’, ‘PI92569’, ‘CItr4174’, and ‘PI43355’ (**Supplementary Table 1 & 2**).

### Genotyping

The NAM population genotyping, curation methods, and population maps were previously reported (Sandhu et al. 2021a, c). The initial genotypic data used in this study composed of 73,345 high-quality markers which were anchored to the Chinese Spring RefSeqv1 reference map (Marcussen et al. 2014) and NAM_175_ selected for stable GPC were used. Individual RIL with missing GPC data were removed before filtering. Markers with more than 20% of data missing were removed for further analysis. Individual RIL missing more than 10% allele data were removed before culling based on markers with less than 10% minor allele frequency (MAF) to lower the probability of Type I error during GWAS. After filtering, 175 RILs from NAM_175_ and 38,588 markers were used for GWAS analysis.

### Population structure and genome-wide association studies

Kinship matrix and structure parameters were calculated by GAPIT (Lipka et al. 2012). The VanRaden algorithm was used to derive the kinship parameter from marker genotypes (VanRaden 2008). Population structure was analyzed using principal component analysis (PCA) implemented with the R function ‘prcomp’ in the software GAPIT (Price et al. 2006). Genetic relatedness among NAM_175_ and NAM_650_ was calculated using population differentiation coefficient (*F*_*st*_) (SAS Institute Inc. 2011). The GWAS was conducted in the GAPIT R package using Bayesian information and Linkage-disequilibrium Iteratively Nested Keyway (BLINK) model (Huang et al. 2018). Association studies were performed on the NAM_175_ using GPC stability index and environmentally-stable phenotypic GPC derived from the FW regression. Selection of PCA groups for GWAS was performed by evaluating Q-Q plot results for each model. PCA groups were included as covariates in the BLINK model to account for population structure. The Bonferroni correction with a stringent α=0.05 was used to identify highly significant associations.

### Genomic selection

Genome-wide marker effects for GPC and GPC stability were estimated using ridge regression best linear unbiased prediction (rrBLUP; Endelman 2011), according to the model:

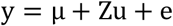

where y is an N x 1 vector of BLUEs for GPC or GPC stability for each RIL, µ is the overall mean, Z is an N x M matrix of markers, u is a vector of marker effects, and e is a vector of residuals. GS was performed with five-fold cross-validation by including 80% of the samples in the training population and predicting the GEBVs of the remaining 20% of the samples under each environmental condition. For accuracy assessment, two 50 replication sets were performed, where each replication consisted of five model iterations.

Genomic selection models were developed for the NAM population of 650 RILs (NAM_650_) and separately for the 175 RILs (NAM_175_) from seven families selected based on lower variation for GPC across environments. Furthermore, independent validations were performed using both sets of RILs for predicting GEBVs for GPC. During independent validations, GS models were trained on the previous year data set, and predictions were made for the upcoming year. Models trained on 2014 GPC data were used for predictions in 2015 and 2016. Similarly, the 2015 GPC training model was used for 2016 predictions (Sandhu et al. 2021c).

## Results

### Variation of grain protein content across environments

The GPC values of the NAM_650_ population ranged from 11.2-18.0% in 2014, 8.7-16.8% in 2015 and 9.7-17.0% in 2016 (**Figure 1**), with 2014 having the highest mean GPC. Distributions of GPC values were normal in the three environments (**Supplementary Figure 1**) with significant positive correlations (R^2^=0.49 for 2014 and 2015, R^2^=0.42 for 2014 and 2016, and R^2^=0.57 for 2015 and 2016). Broad-sense heritability for GPC was moderate to high, ranging between *H*^*2*^=0.62 in 2014, *H*^*2*^=0.36 in 2015 and *H*^*2*^=0.68 in 2016. The average GPC for the 26 families evaluated in this study is presented in **Supplementary Table 1**. Using this data, seven families totaling 175 RILs (NAM_175_) were selected for low variation in GPC (**Supplementary Table 2**).

**Figure 1:**
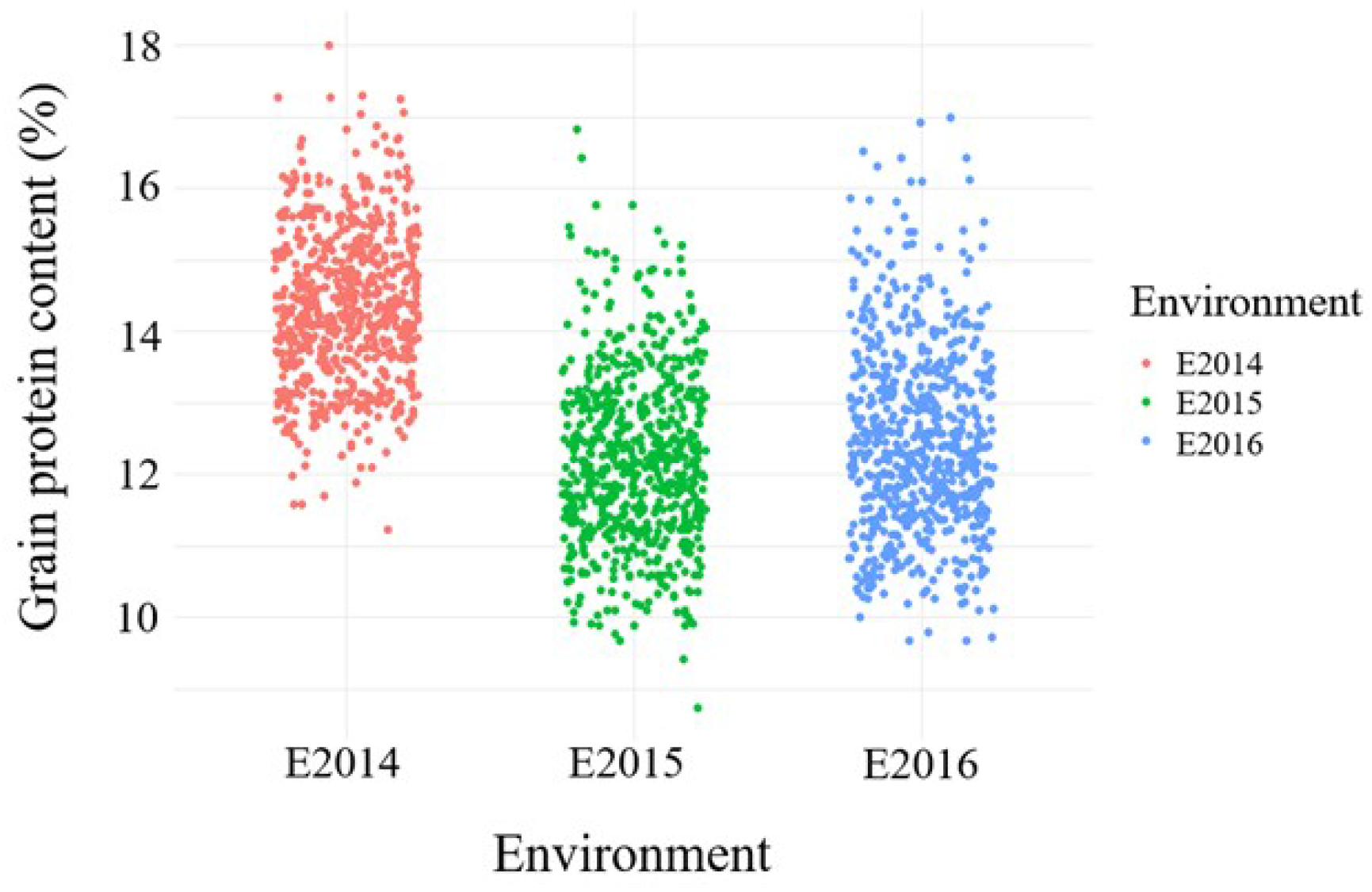
Variation of grain protein content for all the NAM_650_ across the three environments. The X-axis depicts the three environments, namely 2014, 2015, and 2016; the Y-axis shows the GPC in percentage.

### Stability analysis

Stability index (b), environmental effect (t), and genotypic effect (g) values were obtained for the NAM_175_ population. Absolute values of stability index for NAM_175_ RILs ranged from 0.00 to 2.504 and were normally distributed. Thirty RILs were identified which had no significant difference of stability index from 0 using individual t test (*p* < 0.05). Environment 2014 was observed to be the most favorable for high GPC. NAM_175_ population was divided into five categories based upon the GPC and stability index, the trends of which are depicted in **Figure 2**. Categories 2, 3, and 5 includes the stable GPC lines with GPC in the range of 15-17%, 14-15%, and 9-14%, respectively (**Figure 2**). Categories 1 and 4 represent the plastic lines with reversal of GPC when moving to other environments. Categories 1, 2, 3, 4, and 5 includes 81, 15, 9, 64, and 6 RILs, respectively.

**Figure 2:**
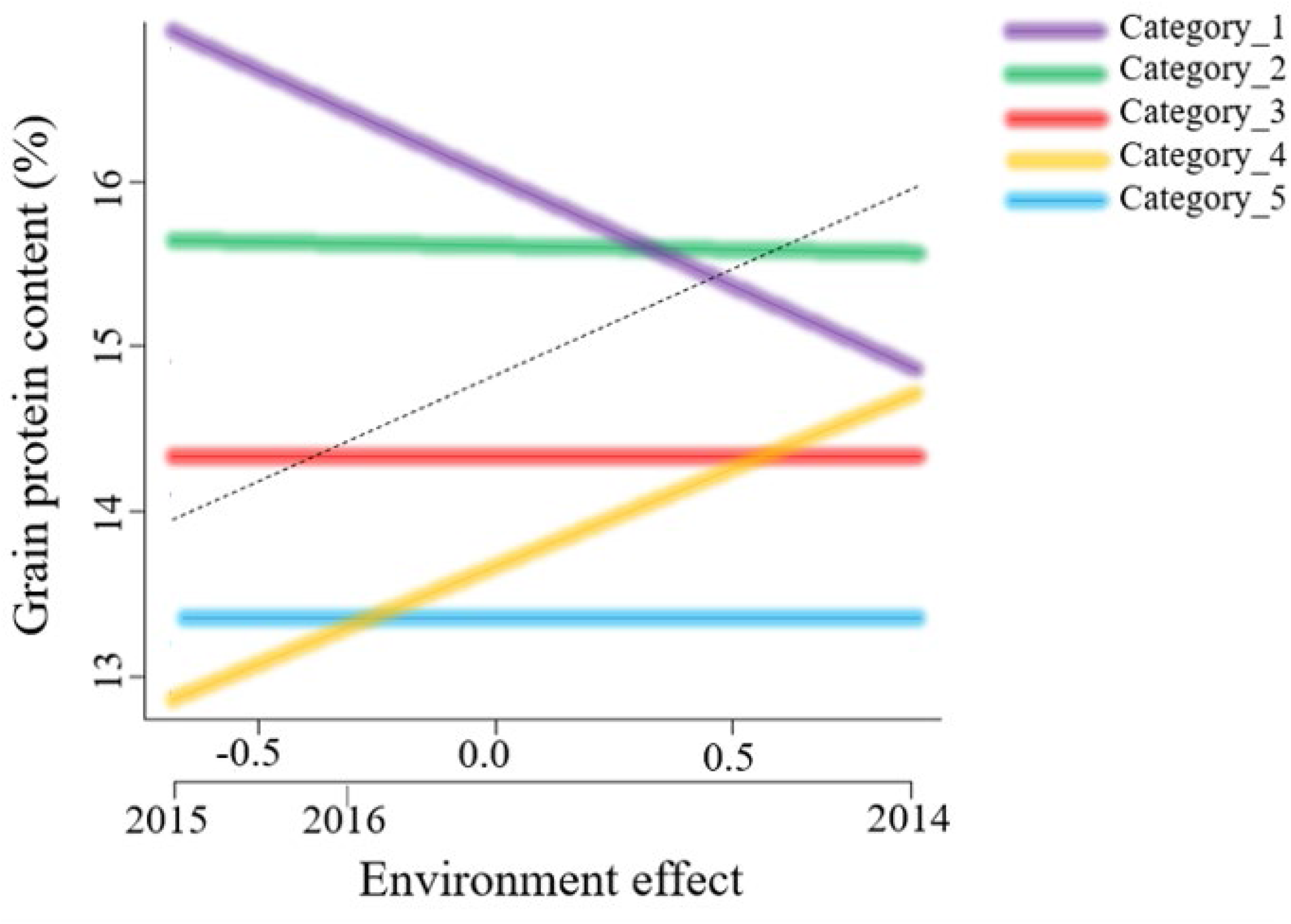
Trends for stability index and GPC observed in the NAM_175_ population. The slopes represent the stability index, X-axis depicts the environmental effect, and Y-axis shows the grain protein content.

### Population structure analysis

Population structure analyzed with PCA separated the NAM_175_ population into seven separate groups based on their different parents, in addition to the common parent ‘Berkut’ (**Figure 3**). The PC1 accounted for 7.0% of the variation, whereas the PC2 explained 5.0% of the genetic variation (**Supplementary Figure 2**). Kinship plots obtained from VanRaden algorithms in GAPIT also separated the population into seven different groups (data not shown). Inclusion of different number of PCs as covariate in the GWAS model demonstrated that the first three PCs best control the false positives and false negatives, as evident from the Q-Q plots (**Supplementary Figure 4)**. *F*_*st*_ coefficient was 0.11 for the NAM_175_ and 0.18 for that NAM_650_.

**Figure 3:**
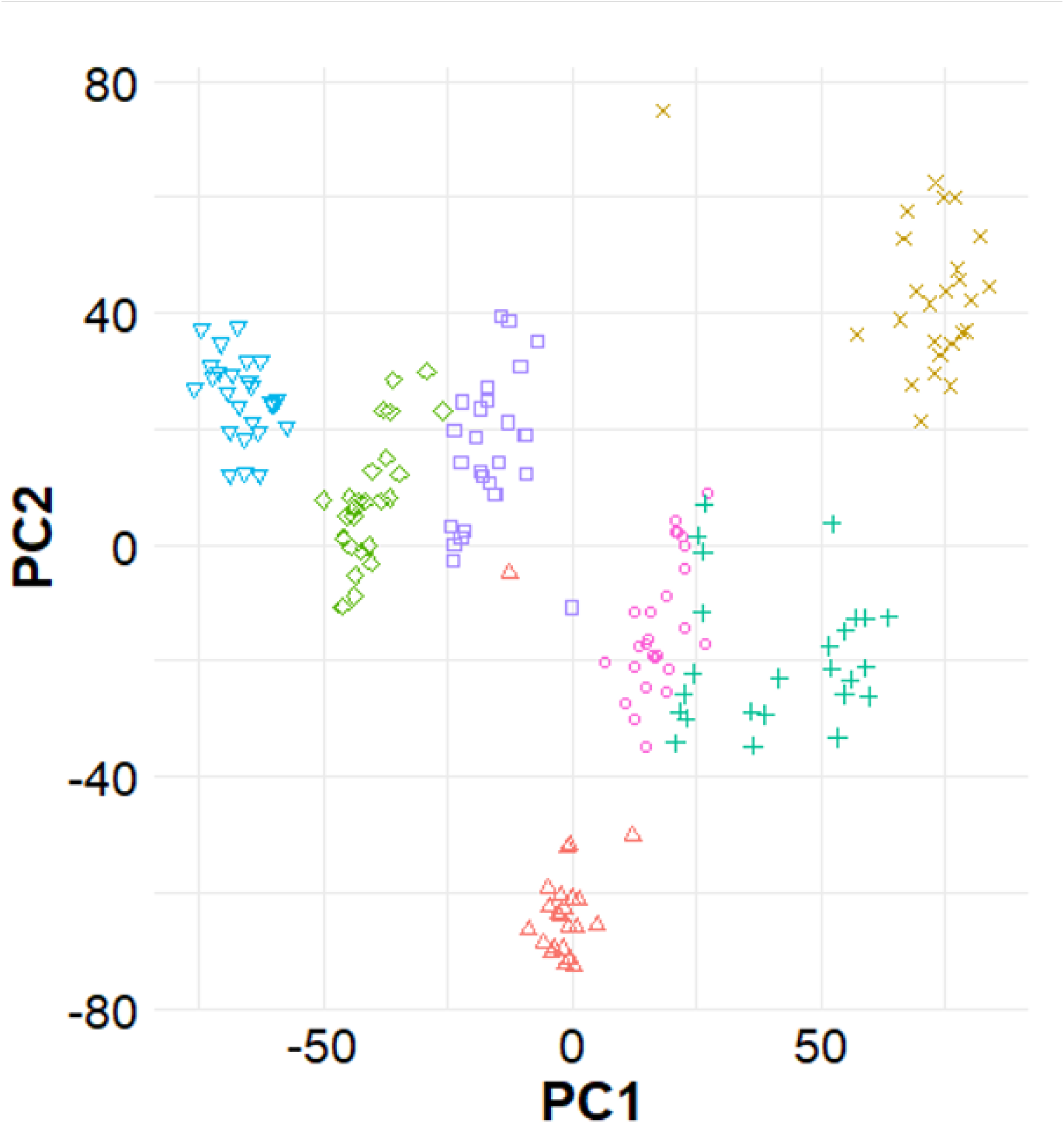
Population structure inferred from principal component analysis and illustrated with the first two principal component’s help. The whole population is subgrouped into seven groups, each group having a single NAM family.

### Marker-trait associations for the stability of grain protein content

Seven QTLs controlling GPC stability were identified on six chromosomes using a stringent Bonferroni correction of α=0.05 (**Table 1; Supplementary Figure 3**). The variation explained by each locus ranged from −4.19 to 3.68%. Two significant QTLs were on chromosome 3B and chromosomes 1A, 1B, 2A, 2B, and 4A each had one significant association. Cumulatively, these loci explained 18.40% of the phenotypic variation. The QTLs on 1A and 2A have a positive effect on increasing GPC stability. The QTL on 1A has the favorable allele from Berkut and Dharwar Dry, while the QTL on 2A has the favorable allele from the landraces, namely CItr15144, PI210945, PI92001, and Dharwar Dry (**Table 1**). Five other QTLs had a negative effect on the GPC stability, and the parents of origin for associated alleles are provided in **Table 1**. Removal of those alleles from a breeding program by selecting for alternative alleles as these loci will favor the development of lines having increased GPC stability.

**Table 1:**
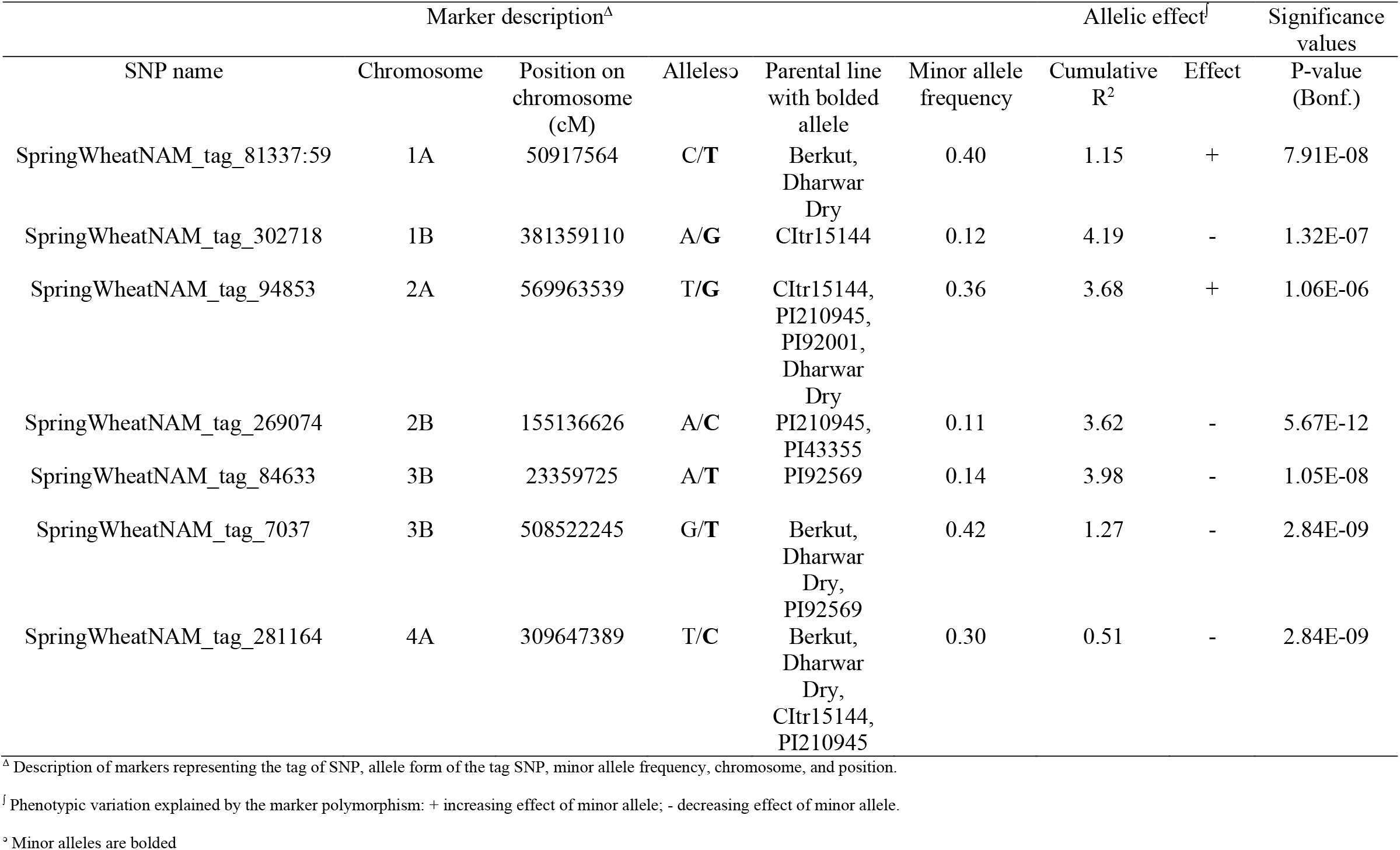
Significant markers representing quantitative trait loci for grain protein content stability in a nested association mapping of hard spring wheat.

Four significant QTLs were identified using the GPC values obtained across the environments as the response trait (**Table 2; Supplementary Figure 5**). GPC was associated with four different loci located on chromosomes 1A, 3B, 4A, and 7A that individually explained −7.30 to 6.78% of the variation for this trait and collectively 16.74% of the total phenotypic variation. The QTL on 1A positively affects GPC, and the associated allele originated from Berkut and Dharwar Dry. The remaining three QTLs negatively affected GPC and removing those alleles will increase GPC. Parents of origin for the associated alleles are provided in **Table 2**. As there are multiple alleles with favorable effects identified from various different parental lines, pre-breeding efforts will be required to introgress all QTL identified. As some QTLs were identified from landraces, selection during this pre-breeding process may help reduce linkage drag.

**Table 2:** Significant markers representing quantitative trait loci for grain protein content in a nested association mapping of hard spring wheat.

| Marker description<sup>Δ</sup> |  |  |  |  |  | Allelic effect<sup>‡</sup> |  | Significance values |
| --- | --- | --- | --- | --- | --- | --- | --- | --- |
| SNP name | Chromosome | Position on chromosome (cM) | Alleles<sup>⊖</sup> | Parental line with bolded allele | Minor allele frequency | Cumulative R<sup>2</sup> | Effect | P-value (Bonf.) |
| SpringWheatNAM_tag_190170 | 1A | 1.24E+08 | T/C | Berkut, Dharwar Dry | 0.49 | 6.78 | + | 1.60E-08 |
| SpringWheatNAM_tag_32264 | 3B | 125543693 | A/T | Berkut, PI43355 | 0.34 | 7.30 | - | 1.26E-06 |
| SpringWheatNAM_tag_82154 | 4A | 6.4E+08 | C/A | CItr15144, PI210945, CItr4175. | 0.26 | 2.34 | - | 1.23E-07 |
| SpringWheatNAM_tag_136322 | 7A | 6.67E+08 | T/G | Berkut, PI43355 | 0.41 | 0.32 | - | 1.55E-08 |

<sup>Δ</sup> Description of markers representing the tag of SNP, allele form of the tag SNP, minor allele frequency, chromosome, and position.
<sup>‡</sup> Phenotypic variation explained by the marker polymorphism: + increasing effect of minor allele; - decreasing effect of minor allele.
<sup>⊖</sup> Minor alleles are bolded

### Prediction for grain protein content and stability

The prediction accuracy for GPC in the NAM_650_ population was *r*=0.50 in 2014, *r*=0.55 in 2015, and *r*=0.53 in 2016. Overall, GPC stability was less predictable with prediction accuracy values between *r*=0.34 and 0.44 and a mean of 0.40. The prediction accuracy of GPC in the selected NAM_175_ subset ranged between *r*=0.56 to 0.69. The maximum prediction accuracy of *r*=0.69 was achieved again for the 2015 environment, while the lowest prediction accuracy of *r*=0.56 was obtained for the 2014 environment. Comparison of prediction accuracies for each environment using the two different sets of population is presented in **Figure 4**. Prediction accuracy is generally high when using the NAM_175_, increasing by 15% as compared to the NAM_650_.

**Figure 4:**
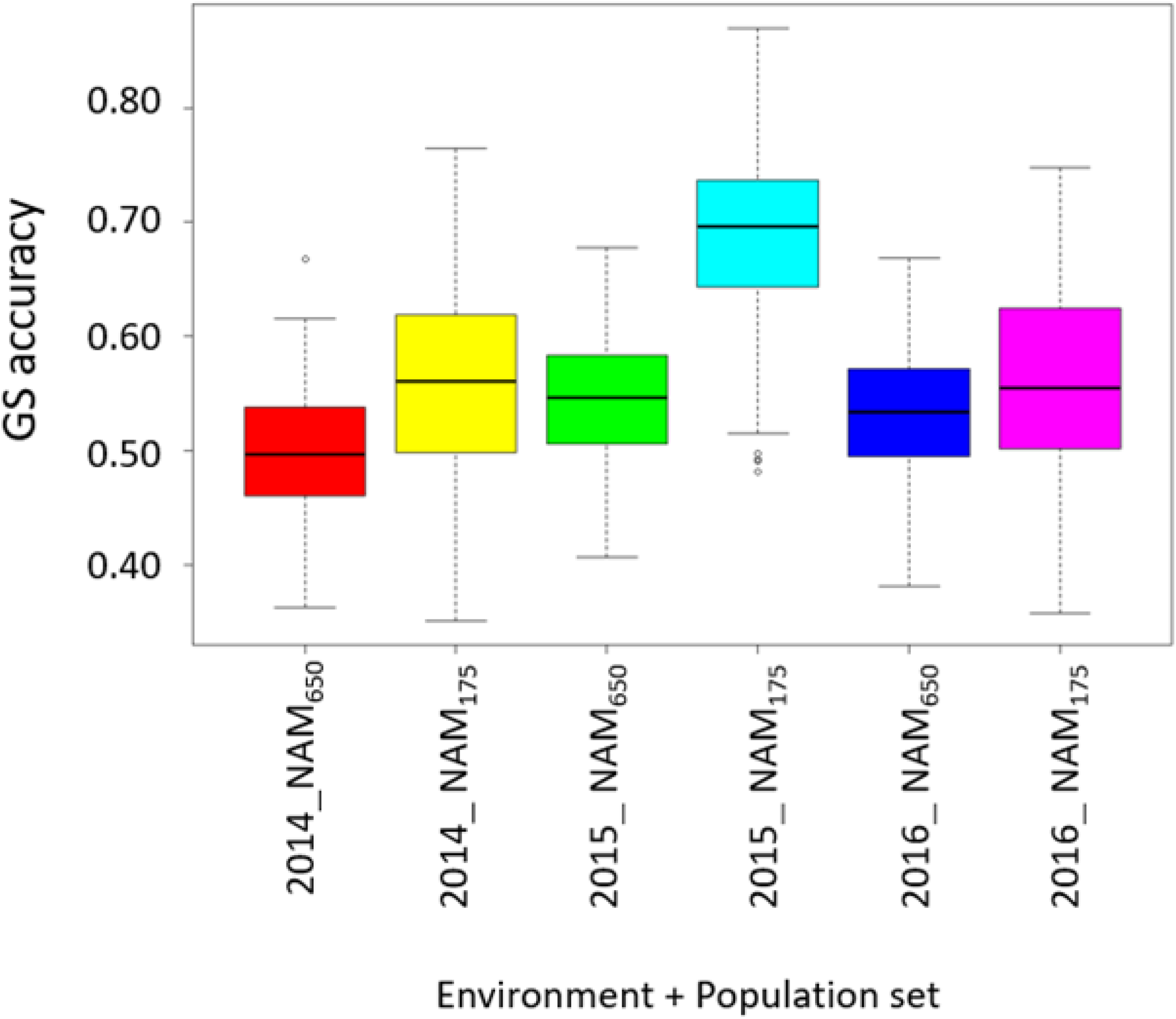
Comparison of GS prediction accuracies for the NAM_650_ and NAM_175_ populations for GPC. The X-axis represents the combination of population and environment, and the Y-axis represents the prediction accuracies.

The GS accuracy was lower for independent validation of each population set and under different environments compared to cross-validation GS accuracies (**Figure 5**). Independent prediction accuracies for the NAM_650_ population ranged between *r*=0.30 to 0.39 and ranged between *r*=0.35 to 0.43 for the NAM_175_ for GPC. The independent prediction accuracies were higher for the NAM_175_, and the same results were observed during the cross-validation prediction scenario. The highest independent prediction accuracy was obtained by a training model on 2015 GPC to 2016 GPC (**Figure 5**).

**Figure 5:**
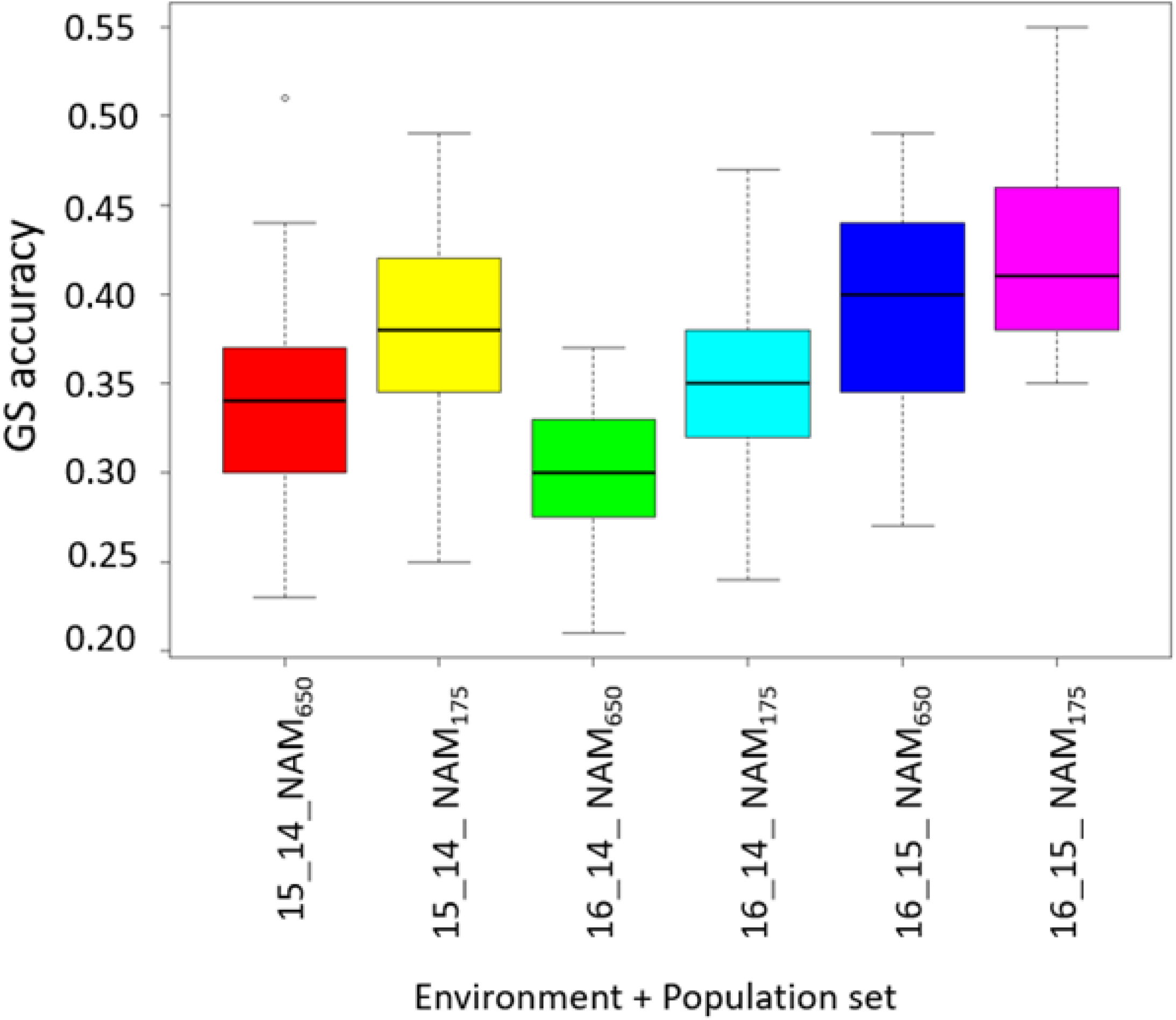
Comparison of independent prediction accuracies for the NAM_650_ and NAM_175_ populations for GPC. The X-axis represents the combination of population and environment where first number represents the testing environment while second number represents the environment on which the GS model was trained, and the Y-axis represents the prediction accuracies.

## Discussion

### Stability of genotypes across environments

A primary goal of plant breeding programs is to select germplasm with superior adaptation to the targeted environments. The performance of genotypes may range from those that are very well adapted to a more narrow set of environments, and perform below average in others, to genotypes that perform consistently relative to others across a wider range of environments and are considered to have greater stability. Herein, we used a static stability concept, targeting a predetermined value of GPC across all the environments. There are numerous statistical tools for analyzing static stability but here we applied FW regression analysis as it is capable of summarizing the interactions in comprehensible ways (Eberhart and Russell 1966).

Crossover and non-crossover interactions were observed in our FW regression analysis which included reversals in rank and scale effects for GPC due to environmental effects and G*E interactions. Identification of only thirty lines with stable GPC across the environments out of 650 lines highlights the challenge of maintaining adequate population size when selecting for a complex quantitative trait in genetically diverse germplasm. Evaluating this population in a greater number of environments would provide more insight into the genetic control of this trait. Additionally, the founder parents of NAM population are primarily landraces that have not undergone routine selection for GPC (Blake et al. 2019) as has been done in contemporary germplasm for at least the past 60 years. The stable lines varied in their average GPC, but lines having high and stable GPC (Category 2) demonstrates the ability to select genotypes for these traits. The lines identified in this study having high GPC and stability are particularly useful for introgression into modern breeding germplasm to expand available allelic variation for this important trait.

### Genomic regions controlling stability of GPC

The QTLs associated with GPC stability and *per se* GPC performance were not detected in the same genomic regions, suggesting that GPC stability and GPC are under the control of different genes. Accounting for the different genetic architecture of these two traits could aid in indexed selection for the desired GPC along with stability. Different QTL controlling yield stability and *per se* yield in wheat has similarly been documented (Berke et al. 1992). In maize, yield and yield stability were observed to be independent, demonstrating the potential for simultaneous selection for both traits (Tollenaar and Lee 2002). Loci controlling yield stability were located in the same regions as QTLs for yield and yield-component in a barley mapping effort (Kraakman et al. 2004). A critical factor that is not captured or quantifiable in each of these yield-related studies are the precise abiotic or biotic factors limiting yield potential in different environments. Thus, stability is an important aspect to consider and our study suggests that breeding programs may select for both GPC and GPC stability by treating them as separate traits for developing varieties for unpredictable environmental variables.

Genomic regions controlling GPC stability have not been investigated in wheat based on available literature. In the present study, seven QTLs controlling GPC stability distributed over six chromosomes were identified that individually explained −4.19 to 3.68% of variation for GPC stability. The loci mapped on 1A and 2A were aiding in increasing the stability of GPC while other five regions were decreasing GPC stability. Similar results were obtained by Sehgal et al. (2017) while mapping genomic regions for yield stability, where 11 QTLs associated with yield stability were distributed on seven chromosomes. The amount of phenotypic variation explained by their yield stability QTLs varied from 3.2 to 8.1%. Thus, although stability is an important consideration when selecting for complex quantitative traits like yield and GPC, appropriate index weighting in genomic selection approaches is most likely the better alternative in selection schemes.

Marker trait associations have reported QTLs for GPC on all the 21 chromosomes of wheat (Blanco et al. 2006; Groos et al. 2003; Joppa et al. 1997; Perretant et al. 2000; Prasad et al. 2003). Loci controlling GPC were mapped to chromosomes 1A, 3B, 4A, and 7A in this study. The locus linked to *SpringWheatNAM_tag_190170* on 1A was previously identified in a DH population using composite interval mapping (Mahjourimajd et al. 2016) and Groos et al. (2003) using an F_7_ RIL population grown in five environments. The favorable allele for this locus was identified from Berkut and Dharwar Dry, both of which are present-day cultivars, suggesting that this favorable locus is fixed in the wheat germplasm and not in the landraces. The 7A GPC locus was discovered in the same region as another investigation (Mahjourimajd et al. 2016; Rapp et al. 2018). Those studies also reported that this locus had a negative effect on the total GPC. Loci on chromosomes 3B and 4A have not been previously reported for GPC in wheat (Blanco et al. 1996; Joppa et al. 1997; Heo and Sherman 2013), which is due to no prior utilization of these landraces in mapping studies (Blake et al. 2019). These two loci had a negative effect on GPC with associated alleles identified in the landraces. The absence of these loci in the cultivars Berkut and Dharwar Dry strengthens the fact that these negative loci have been removed from the germplasm with continuous selection by the breeders. The utilization of diverse landraces in this study provides information about different genomic regions which are not absent in the present-day cultivars.

Similar to GPC stability loci, QTLs controlling GPC *per se* explained a small portion of the variance (16.74%). Given the number of small-effect loci controlling each trait, the GS approach proposed in this study should improve the ability to select for these traits simultaneously during a breeding process. The utilization of indexed genomic selection would assist in selecting simultaneously for GPC stability and GPC (Schulthess et al. 2016). Rapp et al. (2018) demonstrated the use of phenotypic and genomic selection indices to select durum wheat lines for high GPC and grain yield. Similarly, lines having GPC and grain yield were selected using index selection in multi-variate GS model for wheat (Michel et al. 2019). These studies suggested that utilization of index selection in multi-variate GS models would potentially aid in selecting lines having stable GPC in addition to high GPC and grain yield.

### Accuracy for predicting GPC and GPC stability

Prediction accuracy for GPC ranged between 0.50 and 0.69, which is moderately higher than prediction accuracies for grain yield (Battenfield et al. 2016). Heritability of GPC, the effect of the NAM population, and population sizes used for training the GS models would each affect prediction accuracy (Heffner et al. 2011; Poland et al. 2012). Our results are consistent with previous studies where high heritability has resulted in better prediction accuracies in cereals (Windhausen et al. 2012). The heritability of GPC is usually higher than yield, which ultimately resulted in better predictions for GPC (Battenfield et al. 2016). In a genomic prediction experiment for grain yield in oats, lower prediction accuracy was obtained because the experiment was planted under diverse environmental conditions resulting in reduced genetic variance as compared to G*E interactions, and thereby reduced heritability (Asoro et al. 2011). The NAM population in the current study was investigated in more homogenous target environments for wheat production in the PNW. This should result in a relatively lower G*E variance relative to genetic variance, leading to a higher heritability estimate and an increase in genomic selection accuracy.

Prior studies are not available to compare accuracy of GS for GPC stability with stability index values obtained from FW regression in wheat. GS accuracies for GPC stability were significantly (*p* < 0.05) lower than GPC, suggesting a more complex architecture of GPC stability. Huang et al. (2016) conducted GS for grain yield, test weight, and flour protein content stability using an additive main effect and multiplicative interaction (AMMI) model in wheat. They observed an accuracy from 0.14 to 0.31 for the stability of flour protein using four different GS models namely Bayesian ridge regression (0.14), elastic net (0.27), rrBLUP (0.31), and RKHS (0.31). Their study also demonstrated that the stability index for quality traits has less prediction accuracy than the trait itself. Our results, coupled with those of Huang et al. (2016), suggest that GS can be used for predicting GPC stability. Furthermore, the prediction accuracy for GPC stability could be improved by obtaining a stability index from a larger number of field trials, as a large number of environments are useful for more reliable estimations of stability (Piepho 1998). The GS models can be retrained by incorporating data from additional environments in subsequent years. High prediction accuracies for stability could better predict the performance of genotypes at multiple locations during breeding cycles.

There was an improvement of prediction accuracy when model was trained on NAM_175_ compared to the NAM_650_ population. This is in contrast to other studies where it is observed that prediction accuracies improve when the number of individuals in the training population increases (Isidro et al. 2015; Lorenz et al. 2011; Lorenzana and Bernardo 2009). This argument was strengthened by the population differentiation coefficient (*F*_*st*_), suggesting lines in the NAM_175_ were more genetically related compared to the NAM_650_. It has been observed that prediction accuracies increase when training and testing are more genetically related to each other as was the case in NAM_175_ population set (Heffner et al. 2010; Lorenz & Smith 2015). These results suggest that selection for lines having GPC stability also aids in the improvement of GS accuracies for GPC. This study opens up the avenue for utilization of GS in spring wheat breeding program for selecting lines having GPC stability in addition to high GPC.

## Conclusions

Selection for GPC is often secondary to grain yield in terms of breeding objectives for spring wheat, although it is a primary trait producers consider when selecting varieties. We report the first large-scale study of nested association mapping and evaluation of GS for GPC stability in wheat. This study identified wheat lines having less variation for GPC and mapped QTLs controlling GPC stability, an important and often overlooked trait. The identification of stable genotypes with high GPC could help in developing cultivars that can perform similarly in multiple environments. The QTLs identified in this study explained a small amount of phenotypic variation, demonstrating the complexity of the trait which suggests a GS approach could best address breeding for GPC and GPC stability. Prediction accuracy is sufficiently high for the implementation of GS for GPC and GPC stability in this study. With implementation of GS for these traits, predictions can be available while evaluating grain yield, enabling selection for GPC along with agronomic and yield traits.

## Authors contributions

KS: analyzed data, conceptualized the idea, and drafted the manuscript; PM: assisted in data analysis and edited the manuscript; ML: collected the phenotypic data; MP: conducted field trials, and obtained the funding for the project; AC: edited the manuscript, conducted field trials, supervised the study and obtained the funding for the project. All authors read and approved the final manuscript.

